# Ivermectin resistance in dung beetles exposed for multiple generations

**DOI:** 10.1101/2023.05.08.539900

**Authors:** Daniel González-Tokman, Antonio Arellano-Torres, Fernanda Baena-Díaz, Carlos Bustos-Segura, Imelda Martínez M.

## Abstract

Ivermectin is an antiparasitic drug commonly used in cattle, that is excreted in dung, causing lethal and sub-lethal effects on coprophagous non-target fauna. Given that cattle parasites generate resistance to ivermectin, farmers have increased the used doses, with a consequent threat to wild fauna. The dung beetle species *Euoniticellus intermedius* provides ecosystem services by burying dung in cattle pastures, however it is highly threatened by ivermectin. Here we experimentally tested whether *E. intermedius* generates resistance against ivermectin after being exposed for several generations to a sublethal dose. We generated two laboratory lines where beetles were exposed to either ivermectin-treated or ivermectin-free dung for 18 generations. We compared reproductive success (total brood balls, emerged beetles, proportion emerged and days to emergence) of beetles from both lines across generations. Additionally, for each line, we carried-out toxicity experiments with increasing ivermectin concentrations to determine if sensitivity to ivermectin was reduced after some generations of exposure (i. e. if beetles acquired ivermectin resistance by means of transgenerational effects). Our results show that dung beetles do not generate resistance to ivermectin after 18 generations of continuous exposure and quantitative genetic analyses show low genetic variation in response to ivermectin across generations. Together, these results indicate low potential for adaptation to the contaminant in the short term. Although we cannot exclude that adaptation could occur in the long term, our results and comparative evidence in other insects indicate that dung beetles, and probably other species, are at risk of extinction in ivermectin-contaminated pastures unless they are pre-adapted to tolerate high ivermectin concentrations.

## Introduction

Ivermectin is one of the most common antiparasitic drugs used in livestock worldwide (Laing, Gillan & Devaney 2017). It is effective against nematodes and arthropod parasites of humans, cattle and pets and it has even been called a ‘wonder drug’ for its broad spectrum of parasite control and low toxicity for humans (Geary 2005). However, residues of ivermectin are excreted intact in cattle dung and remain active for up to several months in cattle pastures, during which they stay biologically active and threaten non-target coprophagous organisms such as dung flies and beetles (Lumaret *et al*. 2012; Wohde *et al*. 2016). This creates an ecological and economic problem, as coprophagous organisms bury and degrade dung in pastures, helping to maintain soil fertility and eliminating noxious fauna that otherwise would cause livestock disease (Nichols *et al*. 2007). In addition, the economic value of dung beetles in cattle pastures is calculated in up to $423 USD per cow and, therefore, their conservation is urgent to preserve their ecosystem services (Lopez-Collado *et al*. 2017).

Ivermectin in dung reduces the emergence of dung flies and beetles and the most susceptible stages are larvae rather than adults (Lumaret *et al*. 2012). Ivermectin use can be the main threat (besides habitat loss) for dung beetle diversity in cattle pastures, even more than the intensity of farming or the degree of forest fragmentation in the surrounding landscape (Alvarado *et al*. 2017). Ivermectin-treated insects, particularly dung flies and beetles, produce less offspring (Lumaret *et al*. 2012; Blanckenhorn *et al*. 2013; González-Tokman *et al*. 2017) and offspring with reproductive disadvantages such as smaller body size or reduced sexual traits (González-Tokman et al. 2017; Baena-Díaz *et al*. 2018). As ivermectin is slowly excreted in treated cattle, low doses have sublethal effects on the physiology and fitness of dung feeding insects (Verdú *et al*. 2015; González-Tokman *et al*. 2017; Martínez et al. 2017) that even persist across generations (Baena-Díaz *et al*. 2018; Conforti *et al*. 2018). This issue gets more challenging as ivermectin resistance has been reported for several parasites, including nematodes (Shoop 1993; Dent *et al*. 2000; Terrill *et al*. 2001; Kaplan 2004; Osei-Atweneboana *et al*. 2011), mites (Currie *et al*. 2004; Perez-Cogollo *et al*. 2010; Castro-Janer *et al*. 2011; Rodríguez-Vivas *et al*. 2014) and insects (Byford *et al*. 1999), leading farmers to increase the used doses to control livestock parasites.

In arthropods, ivermectin resistance has been observed in some parasitic mites such as *Boophilus microplus* (Benavides & Romero 2000), *Sarcoptes scabiei* (Currie *et al*. 2004; Terada *et al*. 2010) and *Rhipicephalus microplus* (Perez-Cogollo *et al*. 2010). In insects, the evidence of ivermectin resistance is scarce and limited to hematophagous parasitic horn flies (*Haematobia irritans*), that become ca. 3-fold resistant after 23 generations and 6-fold resistant after 50 generations (Byford *et al*. 1999). In *Drosophila melanogaster* flies and *Aedes aegyptii* mosquitoes, resistance to ivermectin is achieved after exposure to other insecticides, revealing cross-resistance (Kane *et al*. 2000; Deus *et al*. 2012). Despite ivermectin resistance occurs, it seems to take longer and be less effective than resistance to insecticides or other antiparasitic drugs, as in *Haematobia irritans* flies, where the magnitude of the resistance was 3-fold with ivermectin to 1470-fold with the insecticide permethrin (Byford *et al*. 1999), probably because of the different physiological mechanisms involved in resistance against different drugs (Kane *et al*. 2000; Seaman *et al*. 2015).

Here we tested for the possibility that dung beetles also generate resistance to ivermectin after being exposed for several generations. To evaluate this idea, we performed an experiment where we exposed a line of beetles to a moderate concentration of ivermectin during 18 generations. In parallel, we grew a control line of beetles that was maintained free of ivermectin for 18 generations. Across generations, we performed toxicity experiments in both lines to test for the effect of increasing ivermectin concentrations on offspring emergence and developmental time. Toxicity experiments allowed to calculate the lethal concentration 50 (LC50) of ivermectin in both lines. By controlling for genetic relatedness between experimental beetles, we also estimated heritability and genetic variation in ivermectin resistance. We predicted: (1) that beetles in the ivermectin exposed line would tend to increase fitness in contaminated dung across generations; (2) that beetles in the ivermectin exposed line, compared to the control line, would show better performance when exposed to increasing ivermectin concentrations, (3) that resistance ratios would increase in the ivermectin exposed line and (4) that there are genetic variation and heritability in ivermectin responses. If these predictions are met, they would indicate that beetles generate resistance to ivermectin after several generations of exposure, giving promising insights regarding parasite management in contaminated pastures. Otherwise, the use of ivermectin would condemn the studied dung beetles to disappear from contaminated pastures.

## Materials and methods

The present study was carried out with the dung beetle *Euoniticellus intermedius* (Coleoptera: Scarabaeinae), which is one of the most fecund species of its subfamily, with a relatively short generation time of ca. four weeks (Martínez et al. 2019). This beetle is native from Africa but was introduced to remove dung from cattle pastures in the United States in the 1970’s and has migrated southwards; despite not being reported as invasive (Del Val et al. 2017), now it is one of the most abundant species in Mexican cattle pastures (Montes de Oca & Halffter 1998) and is particularly threatened by ivermectin since it shows attraction for contaminated dung (Holter, Sommer & Gronvold 1993).

Beetles were collected in San Román ranch, Medellín, Veracruz, Mexico (18°58’19.37” N, 96°04’51.43’’ W; 42 asl) in July 2017. The owners of the ranch report that they do not use ivermectin to control cattle parasites. To start the experiment, we collected 151 females and 100 males and transported them to the laboratory, where the rest of the study was carried-out under insectary conditions (27 ± 1.8 °C; 80% mean humidity). For logistic reasons, beetles were fed cattle dung collected in Palo Alto ranch, Acajete, Veracruz, Mexico (19°35’29.1” N, 97°00’05.5”W), where ranch owners also do not use ivermectin. Before feeding the beetles, dung (80-82% humidity) was frozen for at least 48 h at -22°C to eliminate parasites. Collected beetles were reproduced over two weeks in five containers to obtain a first generation of beetles, known to be free of ivermectin for at least one generation.

Starting in the F1, newly emerged beetles were maintained in randomly formed pairs of a male and a female in 1L plastic containers filled with ca. 700 mL moisted, sterilized sifted soil as substrate. The number of used couples (range 13-43 couples per studied line and generation; Figure 2a) depended on the number and timing of beetle emergences. Each pair could reproduce for three weeks (with ivermectin-treated or control dung; see below). After that time the male and the female were removed, and the number of offspring emerged from each container were recorded. We also recorded the number of larvae that did not emerge from brood balls to have a measurement of female fertility and the time from the pair formation to the emergence of the first offspring, as a measurement of developmental time. To avoid inbreeding, siblings were never crossed with each other. Pairs where the male or the female died before a week were not considered for the analyses.

### Experimental lines

In this experiment (Figure 1) we generated two lines of beetles, one that developed 18 generations in ivermectin (IVM line) and a parallel, not-exposed line (Control line), that developed free of ivermectin during the same 18 generations. Ivermectin acts on invertebrate cell membranes, specifically in glutamate-gated chloride channels, increasing permeability to chloride ions, leading to cell hyperpolarization (Kane *et al*. 2000; Meyers *et al*. 2015). Given that it acts in neurons and muscular cells, it causes paralysis, inhibition of feeding and reproduction, and death (Laing *et al*. 2017).

**Figure 1.**
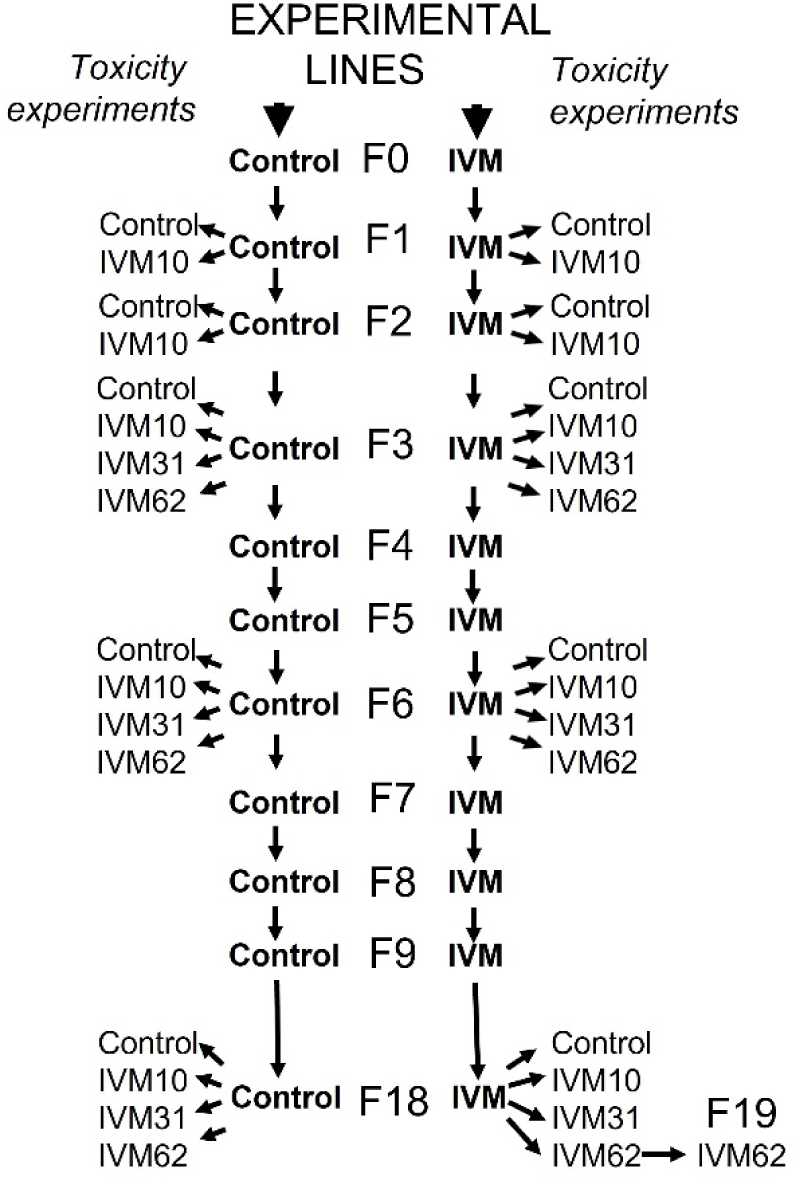
Experimental design to test for ivermectin resistance in *Euoniticellus intermedius* dung beetles across multiple generations of exposure. Field-caught beetles were reproduced in the laboratory for one generation in ivermectin-free dung before starting two experimental lines (F0), one exposed to ivermectin (IVM line, 10 µg of ivermectin per kg of fresh dung) and the free of ivermectin (Control line) until F18. In generations F1, F2, F3, F6 and F18 we performed toxicity experiments to evaluate dung beetle performance at different ivermectin concentrations (IVM10, IVM31 and IVM62, corresponding to 10, 31 and 62 µg of ivermectin per kg of fresh dung). For each line, generation and toxicity experiment (except for the selection lines in the F6) we quantified the number of brood masses produced per couple, the number of emerged beetles, the proportion of emerged beetles and days to emergence. In F18, five couples of emerged beetles from treatment IVM62 were exposed to IVM62 treatment to evaluate the same variables. Sample sizes are given in Figure 2.

**Figure 2.**
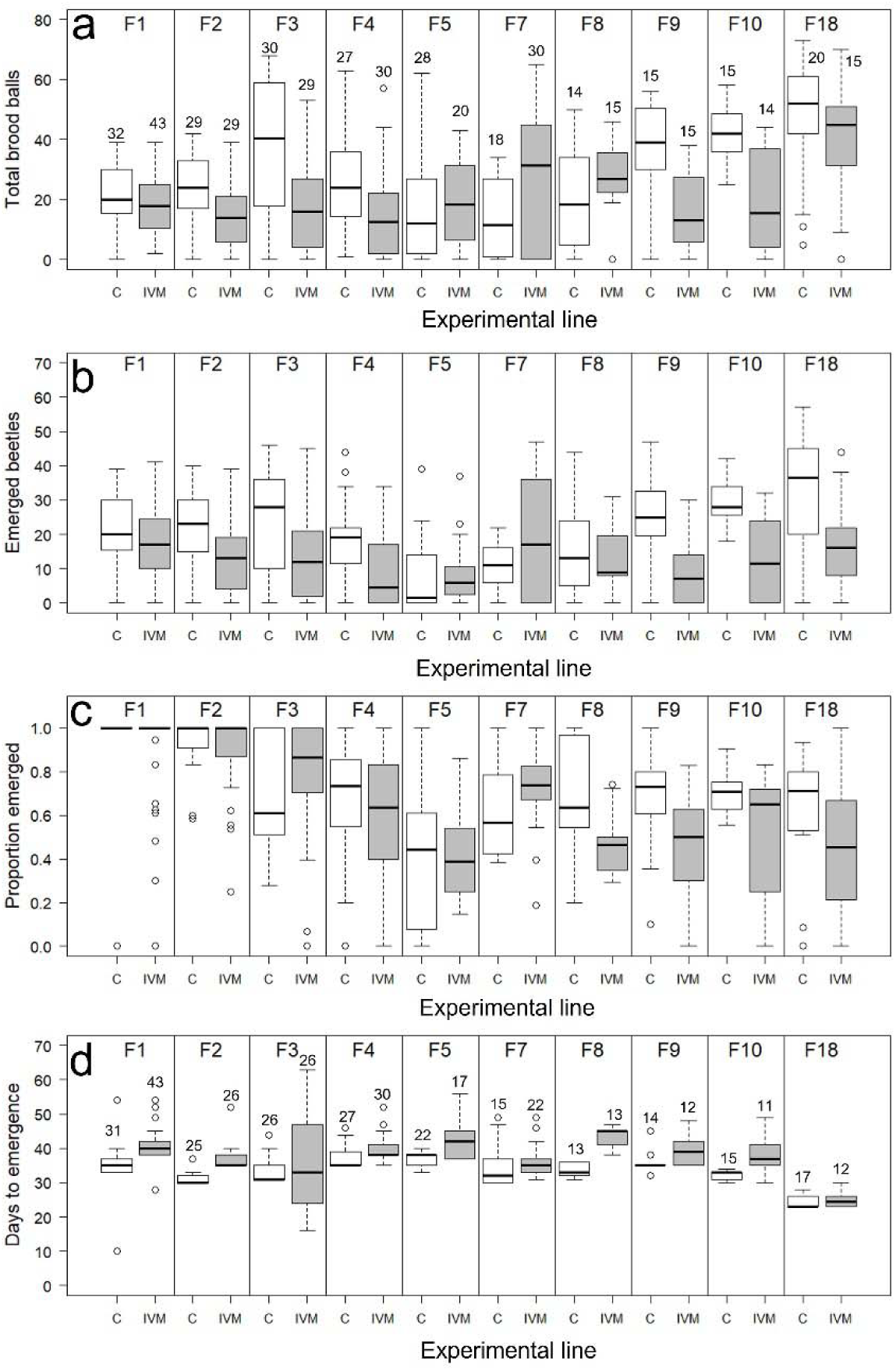
Effect of experimental line (Control or IVM) across generations in *Euoniticellus intermedius* dung beetles. Sample sizes are the same for figures a, b and c and get reduced in the analysis of days to emergence, as there were nests where no beetles emerged (figure d). Numbers next to the bars represent the numbers of analyzed couples.

The experimental treatments were spiked in defrosted dung, which was provided to the beetles three times per week (see similar procedures in (Blanckenhorn *et al*. 2013)). In the treated line (IVM line), beetles from F1-F18 were exposed to ivermectin in the dung (10 µg of ivermectin per kg of fresh dung; Ivermectin, CAS-Number 70288-86-7 Purity of IZ90% ivermectin B1a and IZ5% ivermectin B1b, Sigma-18898). Given that ivermectin was diluted in 50 mL acetone per kg of dung, acetone (CAS-Number 1567-89-1; Sigma purity >99.8%) was used as treatment in the Control line (50 mL per kg of fresh dung). The used ivermectin dose in the IVM line was chosen for being realistic, as it falls in the range of ivermectin excreted by treated cattle after four weeks (Wohde et al. 2016). This dose is considered moderate, as in some populations of *E. intermedius* it has shown to reduce emergence by 50% (Baena-Díaz *et al*. 2018) but in other population it did not affect beetle emergence or physiological condition (González-Tokman *et al*. 2017). As expected, in the present study, the treatment used in the IVM line acted as a moderate selection pressure (see results). In generations F6 and F11-F17 we were not able to register emerged beetles in the IVM lines, so these generations were not considered for statistical analyses, although emerged individuals were used to form the subsequent generations. Generations F11-F17 were maintained in three large terraria per line, containing 20 couples per line but we were not able to monitor the reproductive success in experimental lines, so we just maintained the IVM lines without registering the number of brood balls or emerged beetles. Unexpectedly, in F13 high mortality in both lines left only 12 couples in the IVM line. The Control line in F13 suffered even higher mortality leaving only four females and a male. Therefore, we put together the laboratory population of this particular line with 10 new males and 10 females that had spent a generation in the insectarium feeding control dung in a large terrarium. Both lines got recovered the next generation and 20 couples were formed again for each line. This did not cause any evident effect in the next (and last) evaluated generation (F18), where the Control line maintained similar trends in offspring emergency as past generations (see results).

### Toxicity experiments

From a subset of beetles emerged from both lines (in F1, F2, F3, F6 and F18), we carried out toxicity experiments to evaluate the effect of increasing concentrations of ivermectin (Figure 1). By doing this, we could determine whether individuals from the IVM line (compared to the Control line) became resistant to ivermectin across generations. In F1 and F2, the toxicity experiment consisted of two treatments: ivermectin (10 µg of ivermectin per kg of fresh dung) and control (acetone). In F3, F6 and F18, the toxicity experiment consisted in four treatments with increasing concentrations of ivermectin (10, 31 and 62 µg of ivermectin per kg of fresh dung) plus a control treatment (acetone). These new concentrations (IVM31 and IVM62) are considered high, as they reduce emergence of *E. intermedius* three to four times (particularly females) and body size and muscular mass in both males and females (González-Tokman *et al*. 2017). Moreover, the used ivermectin treatments represent realistic concentrations found in dung of cattle treated 2-4 weeks earlier with the recommended dose (500 µg of ivermectin per kg of cattle body mass (Wohde *et al*. 2016)). As an additional experiment, five couples emerged from IVM62 in F18 were exposed to the same ivermectin concentration (62 µg of ivermectin per kg of fresh dung), but not a single individual emerged in the new generation, which was not considered for statistical analyses. Sample sizes for each generation, line and treatment are shown in Figure 3a. Again, when the male or the female died before a week, the pair was excluded from the analyses. We also estimated the broad sense heritability of reproductive traits using parent-offspring regressions.

**Figure 3.**
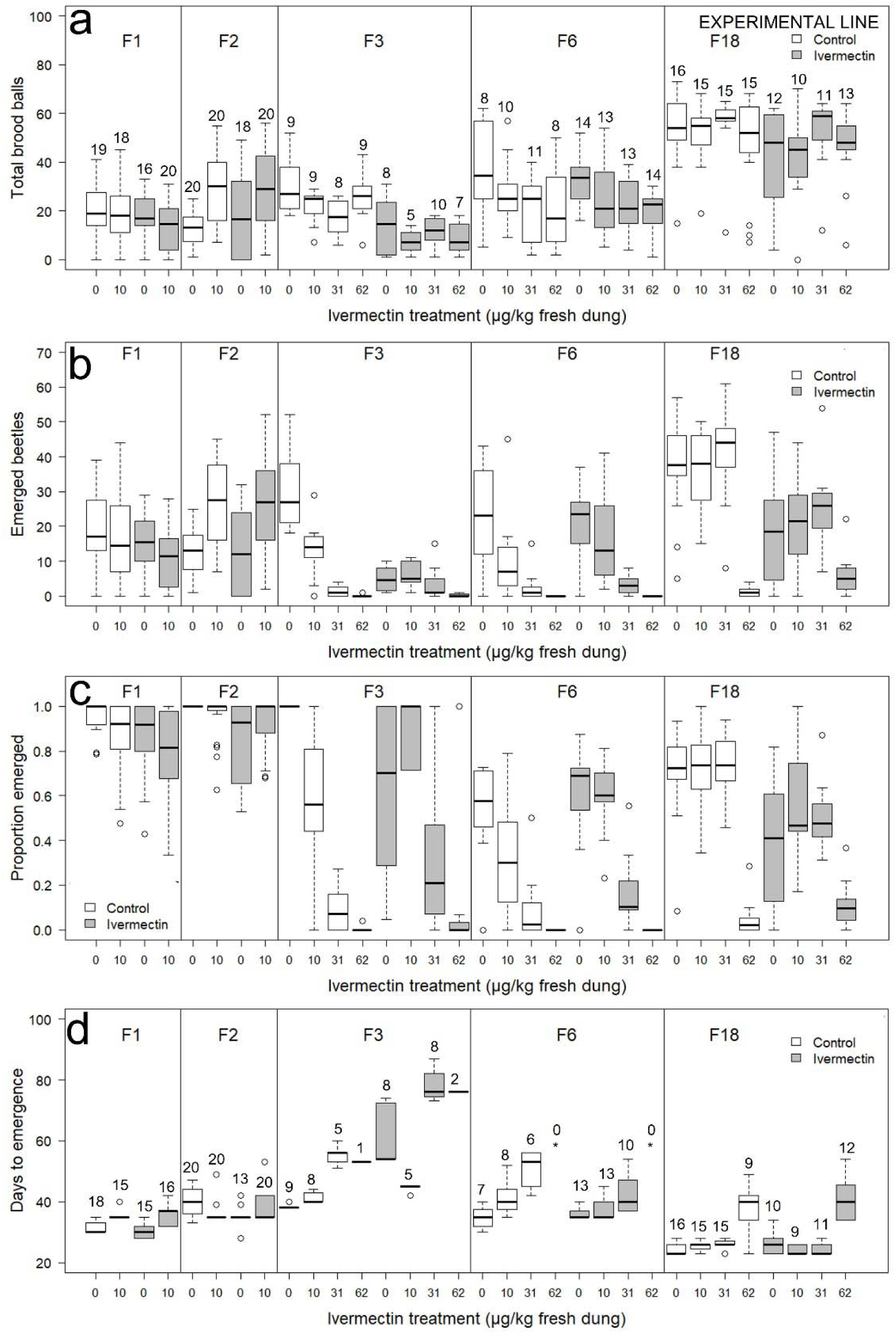
Effect of experimental line (Control or IVM) and ivermectin treatment (0 [control], 10, 31 and 62 µg of ivermectin per kg of fresh dung) across generations in *Euoniticellus intermedius* dung beetles from toxicity experiments. Sample sizes are the same for figures a, b and c and get reduced in the analysis of days to emergence, as there were nests where no beetles emerged (figure d). Numbers next to the bars represent the numbers of analyzed couples. *Represents treatments where there were no emerged beetles and were not considered for the analyses of days to emergence.

### Statistical analyses

Analyses were done according to (Zuur *et al*. 2009; Crawley 2013) in R program (R Development Core Team 2015). To compare the effect of treatment (Control or IVM) across generations, we carried out generalized linear models (GLM) to analyze the total number of brood balls, the number of emerged beetles, the proportion of emerged beetles and the developmental time. For doing so, the used statistical models included the following predictors as factors: Generation, Line and the interaction Generation X Line. The number of brood balls and the number of emerged beetles were analyzed with a GLM with negative binomial errors (given the high overdispersion found for the models with Poisson errors). The proportion of emerged beetles (number of emerged beetles / total number of brood balls) was analyzed with a GLM with quasibinomial errors (given the high overdispersion found for the model with binomial errors). Differences in the number of days to the first emergence were analyzed with a Cox proportional hazards model.

In the toxicity experiments, where different concentrations of ivermectin were tested in F1, F2, F3, F6 and F18, the statistical models also tested the effect of treatment and the triple interaction Generation X Line X Treatment. Given that the triple interaction was significant in most analyses (Table 2), we carried out separate analyses for each generation. These new analyses initially tested the effect of Line, Treatment and the interaction Line X Treatment. The original models were reduced based on the Akaike Information Criterion (AIC, for the case of total number of brood balls and number of emerged beetles) and step by step (removing non-significant terms) for the proportion of emerged beetles and time to the first emergence (as AIC is not available for quasibinomial GLMs or Cox proportional hazards models).

In toxicity experiments of F3, F6 and F18, where we tested several ivermectin concentrations, we estimated ivermectin LC50 for Control and IVM lines with logit analyses in R package ecotox (Hlina 2020). Resistance ratios (RR), were estimated in F3, F6 and F18 as LC50 in the IVM line / LC50 in the Control line (Mazzarri & Georghiou 1995). Values of RR larger than 3 are considered resistance and values from 1.5-3 are considered tolerance rather than resistance (Byford *et al*. 1999).

We performed parent-offspring regressions to estimate the broad sense heritability of number of brood balls, number of emerged beetles, proportion of emerged beetles and days to first emergence. We performed separate regressions for daughters and sons, using the values of each couple as the parental value (explanatory variable) and the values of the respective couple for daughters and sons as the offspring values (response variable). The coefficient estimate for the parental value was taken as the broad sense heritability (Falconer & Mackay 1996) and its standard error was used to calculate the statistical significance with a z ratio test. A positive significant slope would indicate a significant contribution of genetic variation to the total phenotypic variation of each trait. The regressions were run using a linear model with normal error distribution, including line and generation as covariates. The interactions between line and parental traits were also tested but excluded from final models since they did not explain the observed variation. The proportion of emerged beetles was logit transformed to improve normality.

## Results

*Euoniticellus intermedius* dung beetles did not improve performance in ivermectin-contaminated dung after being exposed for 18 generations to a moderate concentration of the contaminant (Figures 2 and 3). The effects of Line, Generation and the interaction Line X Generation were significant for most analyzed variables (Table 1), but beetles from the exposed line (IVM) did not improve performance (mainly number of emerged beetles but also total brood balls, proportion of emerged beetles and developmental time) in contaminated dung across generations (Figure 2). Moreover, in the last three monitored generations (F9, F10 and F18), the negative effect of the experimental line on the number of emerged beetles was more evident than in earlier generations (Figure 2b).

**Table 1.**
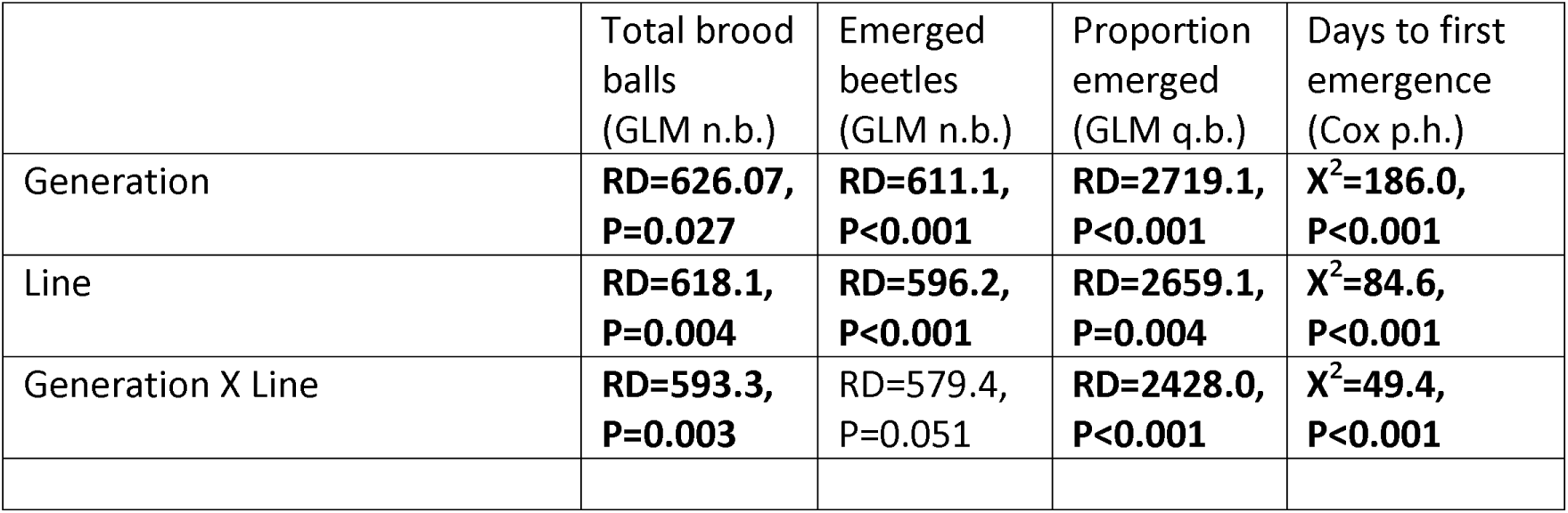
Effect of experimental line (Control or IVM) in *Euoniticellus intermedius* dung beetles across generations. GLM n.b.=negative binomial GLM; GLM q.b.=quasibinomial GLM; Cox p.h.=Cox proportional hazard regression; RD=Residual deviance; Χ^2^=Chi-squared. Significant effects are shown in bold.

|  | Total brood balls<br>(GLM n.b.) | Emerged beetles<br>(GLM n.b.) | Proportion emerged<br>(GLM q.b.) | Days to first emergence<br>(Cox p.h.) |
| --- | --- | --- | --- | --- |
| Generation | **RD=626.07,**<br>**P=0.027** | **RD=611.1,**<br>**P<0.001** | **RD=2719.1,**<br>**P<0.001** | **$\chi^2=186.0,$**<br>**P<0.001** |
| Line | **RD=618.1,**<br>**P=0.004** | **RD=596.2,**<br>**P<0.001** | **RD=2659.1,**<br>**P=0.004** | **$\chi^2=84.6,$**<br>**P<0.001** |
| Generation X Line | **RD=593.3,**<br>**P=0.003** | RD=579.4,<br>P=0.051 | **RD=2428.0,**<br>**P<0.001** | **$\chi^2=49.4,$**<br>**P<0.001** |

**Table 2.** Effect of experimental line (Control or IVM) and ivermectin treatment (0 [control], 10, 31 and 62 µg of ivermectin per kg of fresh dung) in *Euoniticellus intermedius* dung beetles across generations. GLM n.b.=negative binomial GLM; GLM q.b.=quasibinomial GLM; Cox p.h.=Cox proportional hazard regression; RD=Residual deviance; Χ^2^=Chi-squared. Significant effects are shown in bold.

|  | Total brood balls<br>(GLM n.b.) | Emerged beetles<br>(GLM n.b.) | Proportion emerged<br>(GLM q.b.) | Days to first emergence<br>(Cox p.h.) |
| --- | --- | --- | --- | --- |
| Generation | **RD=573.5,**<br>**P<0.001** | **RD=1037.7,**<br>**P<0.001** | **RD=5421.8,**<br>**P<0.001** | **$\chi^2=182.2,$**<br>**P<0.001** |
| Line | **RD=564.8,**<br>**P=0.003** | **RD=1020.7,**<br>**P<0.001** | **RD=5291.2,**<br>**P<0.001** | **$\chi^2=4.8,$**<br>**P=0.029** |
| Treatment | RD=561.1,<br>P=0.301 | **RD=659.5,**<br>**P<0.001** | **RD=2871.7,**<br>**P<0.001** | **$\chi^2=252.2,$**<br>**P<0.001** |
| Generation X Line | **RD=540.3,**<br>**P<0.001** | **RD=642.0,**<br>**P=0.001** | **RD=2728.4,**<br>**P<0.001** | **$\chi^2=42.1,$**<br>**P<0.001** |
| Generation X Treatment | **RD=502.7,**<br>**P<0.001** | **RD=522.4,**<br>**P<0.001** | **RD=2223.4,**<br>**P<0.001** | **$\chi^2=89.6,$**<br>**P<0.001** |
| Line X Treatment | RD=499.7,<br>P=0.392 | **RD=485.3,**<br>**P<0.001** | **RD=2019.8,**<br>**P<0.001** | $\chi^2=4.8,$<br>P=0.189 |
| Generation X Line X Treatment | RD=496.6,<br>P=0.926 | **RD=488.8,**<br>**P=0.036** | **RD=1790.9,**<br>**P<0.001** | **$\chi^2=38.8,$**<br>**P<0.001** |

Toxicity experiments carried out in generations F1, F2, F3, F6 and F18 confirmed that the negative effect of ivermectin is not reduced after 18 generations. This was observed as beetles from the IVM line did not improve performance (mainly number of emerged beetles but also total brood balls and proportion of emerged beetles) in contaminated dung across generations or when compared to beetles in the Control line (Figure 3; Tables 2 and 3). The significant statistical interaction Line X Treatment (Table 2) showed that differences between beetles from the IVM and control lines changed across generations. However, these interactions did not show a consistent improvement in the performance of beetles from IVM line compared to the Control line in either the same or higher ivermectin concentrations (Figure 3). For example, such a trend was observed in the proportion of emerged beetles in F6 but not in F18 (Table 2; Figure 3c). Also, in F3 the number of emerged beetles in the IVM line was consistently lower than in the Control line. Notably, even in the highest ivermectin concentration (62 µg of ivermectin per kg of fresh dung), beetles from both lines built as many brood balls as those in the lowest concentration (Figure 3a), although such balls rarely emerged (Figure 3c). A different pattern was observed in F18, where the number of emerged beetles was surprisingly high at the highest ivermectin concentration in the IVM line, and was even higher than in the same treatment from the Control line.

**Table 3.**
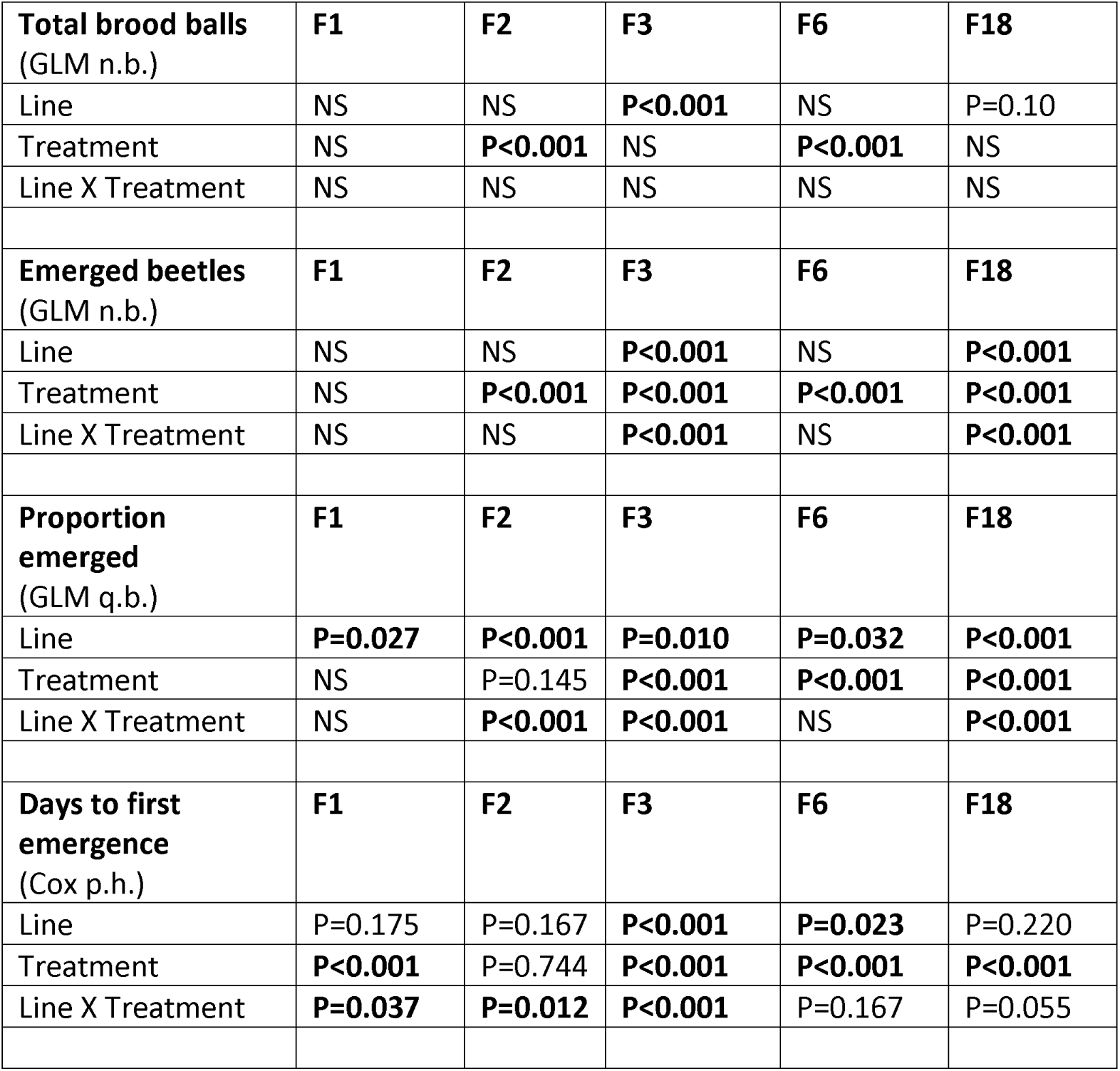
Effect of experimental line (Control or IVM) and ivermectin treatment (0 [control], 10, 31 and 62 µg of ivermectin per kg of fresh dung) across generations in *Euoniticellus intermedius* dung beetles. GLM n.b.=negative binomial GLM; GLM q.b.=quasibinomial GLM; Cox p. h.=Cox proportional hazard regression; RD=Residual deviance. Significant effects are shown in bold.

However, when analyzing ivermectin lethality, ivermectin resistance ratios (RR) indicated lack of resistance and only small tolerance to the contaminant in generation F3, as LC50 of ivermectin in the IVM line was 1.72 times higher than in the Control line (LC50=21.0 versus 12.2 µg of ivermectin per kg of fresh dung; RR=1.72; Figure 3c). However, the RR>1 found in the F3 is mainly explained by the fact that unexposed individuals from the IVM line emerged in very low numbers in this generation (5-6-fold less than unexposed individuals in the Control line) and not because of improved reproductive success at high ivermectin concentrations (Figure 3b). Also, resistance was low in the F6, as the RR was only 1.65 (LC50=13.5 versus 8.2 µg of ivermectin per kg of fresh dung in IVM and Control lines, respectively). In F6, unexposed individuals from both lines emerged in similar numbers but individuals exposed to 10 and 31 µg/kg in the IVM line had higher proportion of emerged offspring than exposed individuals from the Control line, indicating some tolerance. Finally, in the F18, resistance was not evident at all, as RR= 0.56 (LC50=15.2 versus 27.3 µg of ivermectin per kg of fresh dung in IVM and Control lines, respectively). Development times also did not improve significantly in exposed beetles from the IVM line across generations or when compared with the Control line (Tables 1 and 2; Figure 3d).

The parent-offspring regressions were not significant for the number of brood balls, number of emerged beetles, proportion of emerged beetles and days to first emergence, indicating that genetic variation does not explain the phenotypic variance for those traits in the studied dung beetles (Table S1).

## Discussion

In the present study we show that dung beetles *Euoniticellus intermedius* do not improve performance in ivermectin after 18 generations of exposure and that genetic variation does not explain variation in the observed responses, contrary to our four predictions. Beetles growing in ivermectin during 18 generations did not improve reproductive success in contaminated dung across generations. Moreover, the last three studied generations were more severely affected by ivermectin than earlier generations, indicating an amplification of adverse effects of ivermectin across generations on the measured traits. Also, beetles exposed for 18 generations to a low ivermectin dose did not improve performance at higher concentrations, as observed by low resistance ratios, which were even <1 in F18. Therefore, descending from a genetic line that has been exposed to ivermectin for 18 generations not only does not improve performance in contaminated dung, but also may have considerable negative effects in non-contaminated food, as observed by the lower reproductive success in unexposed individuals of IVM than Control line. Our evaluation of ten generations with controlled kinship showed that reproductive traits are hardly heritable, which can explain the observed patterns of lack of resistance. These findings give a pessimistic scenario for dung beetles in ivermectin-contaminated pastures around the world, as ivermectin-treated cows excrete, during the first 28 days post-treatment, contaminated dung with doses that are highly lethal for our studied beetles (i. e. higher than 10 µg of ivermectin per kg of fresh dung) (Wohde *et al*. 2016; González-Tokman *et al*. 2017). It is also plausible that higher doses might increase the selective pressure of ivermectin, facilitating evolution, but this possibility remains to be tested. Our results highlight the need for multigenerational assessments of ivermectin effects in non-target fauna in contaminated pastures.

Ivermectin resistance has been studied in parasitic nematodes, parasitic ticks and only one parasitic insect. In nematodes, three generations are enough to generate resistance to ivermectin (Coles, Rhodes & Wolstenholme 2005) whereas in mites *Sarcoptes scabiei*, ivermectin resistance has been reported after 30 and 58 exposure events (i. e. generations) (Currie *et al*. 2004). In horn flies *Haematobia irritans* (Diptera: Muscidae), the only insects studied for ivermectin resistance, 3-fold resistance is detected after 23 generations and reaches 6-fold after 60 generations (Byford *et al*. 1999). Our study was carried out for 18 generations across 22 months in the laboratory. Even though we cannot discard that resistance could improve after more generations, as shown in horn flies after 23 generations, we did not detect any trend in that direction. Moreover, individuals in the ivermectin-exposed line did not perform better in ivermectin when exposed to moderate and high ivermectin concentrations (31 and 62 µg/kg; Figure 3b). Unlike parasites, which are highly combatted with antiparasitic drugs and therefore they are permanently exposed to these drugs, non-target organisms such as dung beetles may face intermittent exposure to the contaminant, threatening some but not all generations. We also cannot discard that the observed reductions in the number of emerged beetles in some of our studied generations has resulted from potential deleterious effects of ivermectin eroding genetic diversity and causing genetic drift, thus preventing dung beetle adaptation to ivermectin.

Pesticide resistance in insects may be provided by different physiological mechanisms. For example, mutations in glutathione transferases, a family of antioxidant enzymes involved in detoxification and elimination of free radicals, provide insect resistance to DDT, organophosphates and pyrethroids (Enayati, Ranson & Hemingway 2005). In the case of ivermectin, evidence in lice show that lice exposed to a sublethal concentration become more tolerant to a lethal dose later in their lives (Yoon *et al*. 2011); the increased survival is associated to the overexpression of detoxification genes involved in the metabolism of ivermectin. In *Anopheles gambiae* mosquitoes exposed to ivermectin, mechanisms of resistance are associated with the overexpression of immune-response genes (Seaman *et al*. 2015). In the fruit fly *Drosophila melanogaster*, ivermectin resistance is acquired by individuals selected for another antiparasitic drug (nodulisporic acid), and this crossed-resistance is given by glutamate-gated chloride channels (Kane *et al*. 2000). In fruit flies resistant to another macrocyclic lactone, abamectin, resistance is given by overexpression of P-glycoprotein, a transmembrane ATP-dependent drug efflux pump (Luo, Sun & Wu 2013). The extent to which these mechanisms may favor adaptation to ivermectin in dung beetles remains to be studied.

Our quantitative genetic analyses show low genetic variation for ivermectin response, indicating low potential for adaptation to ivermectin in the studied dung beetles, as previously reported in dung flies (González-Tokman et al. 2022). Nevertheless, pesticide resistance can evolve by different means, which we cannot discard. First, standing genetic variation may provide resistance prior to the existence of the pesticide (Hawkins *et al*. 2019), and several generations of exposure could make evident favorable combinations. Further studies in *E. intermedius* populations within its native range, where higher genetic variation is expected, would indicate if some genetic variants and combinations can generate more resistant phenotypes. As a second mechanism of evolution, *de novo* mutations could increase resistance due to random processes (Hawkins *et al*. 2019), and this could be explored in experimental lines exposed to higher mutation rates (i. e. Wendell et al. 2000). The present experimental evidence also shows that phenotypic plasticity and transgenerational effects are not providing any survival or reproductive benefit, as individuals growing up in ivermectin, and their offspring, did not perform better when consistently exposed to the contaminant. This contrasts with previous studies showing high plasticity in response to ivermectin (González-Tokman et al. 2022) and parental effects (Baena-Díaz et al. 2018) affecting subsequent generations of ivermectin-exposed insects. However, the low observed heritability of the measured traits indicates low evolutionary potential in response to ivermectin. Fast evolution could be experimentally evaluated with artificial selection experiments, where only the fittest genotypes contribute to the next generation, or with the use of IVM doses that are higher than the LC50.

Our studied dung beetle, *E. intermedius*, is highly adaptable to new environmental conditions and has colonized several habitats in different continents, probably due to the high female fecundity, high reproductive rate and short developmental time compared to related species of dung beetles (Montes de Oca & Halffter 1998). Even with such high adaptive and invasive potential, this beetle could not improve performance or generate resistance against low doses of ivermectin after 18 generations of exposure in the laboratory. Considering that the study site is dominated by cattle pastures and approximately half of the farmers use ivermectin (González-Gómez et al. 2018), other species of dung beetles with lower reproductive potential and longer developmental times, will hardly become resistant to ivermectin, unless pre-adaptation, standing variation or random mutation provide protection (Hawkins *et al*. 2019). Further studies in other species of dung-degrading organisms, including native dung beetles, are needed to know if some species will develop ivermectin resistance and can still contribute to dung degradation and soil fertilization in ivermectin-contaminated pastures. This is particularly true as ivermectin sensitivity is highly clustered phylogenetically, with different species within a genus varying up to 500 times in sensitivity to ivermectin (Puniamoorthy *et al*. 2014).

The effectiveness of ivermectin has led to use it as a prophylactic treatment applied massively in humans for controlling malaria-transmitting mosquitoes (Alout *et al*. 2014). However, it is of current concern whether these mosquitoes will also generate resistance against ivermectin (Pooda *et al*. 2015). Further studies in target and non-target arthropods are needed to evaluate the genetic and physiological mechanisms of ivermectin resistance and the extent to which different arthropod species generate resistance to ivermectin.

## Acknowledgements

The authors thank Yorleny Gil Pérez, Ana Kory Martínez Diego and Ricardo Madrigal Chavero for logistic and technical support. FBD is supported by a postdoctoral fellowship CONACYT. This study was funded by CONACYT Ciencia Básica project number 257894 granted to DGT.

---

**Table S1.** Regressions coefficients of parental traits on offspring traits (Broad Sense Heritability) across 9 generations of selection. Selection line and generation were included as covariates, and were significant for all the models (except Line for models analyzing Emergence proportion). The interaction between parental traits and line was not significant and excluded from final models. Number within parentheses indicate the S.E. of each estimate. All estimates were not statistically significant with an alpha=0.05.

|  | Total brood balls | Emerged beetles | Emergence proportion (logit) | Days to emergence |
| --- | --- | --- | --- | --- |
| Offspring trait (daughters) | 0.025<br>(0.070) | 0.091<br>(0.074) | 0.057 (0.091) | 0.039 (0.080) |
| Offspring traits (sons) | -0.001<br>(0.072) | -0.038<br>(0.075) | 0.041 (0.10) | 0.039 (0.093) |

